# SbmA coordinates iron homeostasis and antimicrobial susceptibility in *Klebsiella pneumoniae* through phosphorylation-dependent regulation

**DOI:** 10.64898/2026.04.25.720692

**Authors:** Chelsea Reitzel, Jonathan Sayewich, Stevan Cucic, Oscar Romero, Norris Chan, Jennifer Geddes-McAlister

## Abstract

Iron is an essential nutrient that underpins fundamental biological processes, yet its bioavailability is severely restricted during infection due to oxidation and host-mediated sequestration. In Gram-negative pathogens, such as *Klebsiella pneumoniae*, iron limitation imposes a critical selective pressure, necessitating tightly regulated acquisition systems to support growth, virulence, and survival. While canonical pathways for iron uptake are well characterized, regulatory mechanisms coordinating these processes remain incompletely understood. Here, we applied mass spectrometry–based phosphoproteomics to identify iron-responsive regulatory events associated with bacterial iron homeostasis. This approach revealed iron-dependent phosphorylation of SbmA, a conserved inner membrane transporter previously implicated in the uptake of antimicrobial peptides and related substrates. Functional characterization demonstrated that deletion of *sbmA* results in reduced intracellular iron levels and altered cellular morphology, supporting a role in iron acquisition. Complementary proteome mapping of Δ*sbmA* revealed compensatory production of siderophore receptors and TonB-dependent transport systems, further implicating SbmA in maintaining iron balance. Leveraging these findings, integration with high-throughput drug screening identified a compound that exploits SbmA-mediated transport to inhibit bacterial growth, highlighting its potential as a therapeutic entry point. Collectively, this work uncovers a previously unrecognized role for SbmA in iron homeostasis and demonstrates the power of phosphoproteomics to identify condition-specific regulators of essential bacterial pathways. These findings position SbmA as a promising target for antimicrobial development in *K. pneumoniae*.

## Introduction

Iron is an important nutrient metal for nearly all living organisms, serving central roles in diverse biological systems (1, 2). In bacteria, such as Gram-negative *Klebsiella pneumoniae,* iron is one of the most abundant transition metals within the cell, participating in redox reactions to facilitate key biochemical processes, such as cellular respiration, protein/DNA synthesis, and metabolism (3–5). *K. pneumoniae* colonization can lead to severe infections, including pneumonia, urinary tract infections, bloodstream infections, and meningitis, particularly in immunocompromised patients and hospital settings with limited treatment options available (6–9). For this pathogen, iron availability regulates many biological functions, including bacterial growth, virulence, and biofilm formation (10), which are relevant during host invasion. However, within aerobic environments Fe²⁺ is rapidly oxidized to insoluble Fe³⁺, making iron significantly less bioavailable(11), and host cellular processes are designed to scavenge and sequester free iron through production of heme, ferritin, transferrin, lactoferrin, and calprotectin (12). As a result, during infection, free iron is scarce, requiring invading bacteria to adapt and evolve specialized acquisition systems to overcome this nutritional immunity (13, 14).

Ferrous iron (Fe²⁺), which is more soluble under anaerobic or acidic conditions, is thought to cross the outer bacterial membrane through non-specific porins by passive diffusion (15). In contrast, to acquire ferric iron (Fe³⁺), bacteria synthesize siderophores, which are small, high-affinity chelators, to scavenge ferric iron from the environment or host proteins (16). The resulting ferrisiderophore complexes are recognized and transported across the outer-membrane by TonB-dependent transporters (e.g., FepA, FhuA) (17, 18). These transporters rely on the TonB–ExbB–ExbD complex, which harness the proton motive force (PMF) to drive active uptake across the outer membrane (19). Conversely, transport of iron across the inner membrane is mediated by distinct systems depending on the iron species: Feo system for Fe2+, primarily via the GTP-dependent transporter FeoB (20); and ATP-binding cassette (ABC) transporters, such as FhuBC (21) or FepGD (22), for ferrisiderophore complexes. Once inside the bacterial cytoplasm, iron is released from the siderophore and converted to a less reactive and more biologically usable form (23).

Although, many transporters, complexes, and acquisition strategies have been purposed and studied across diverse bacterial pathogens, advances in genome annotation, structural and molecular biology, and predictive modeling continue to reveal previously unrecognized metal ion transport proteins. For example, our previous work explored the influence of iron and zinc availability on *K. pneumoniae* virulence using mass spectrometry-based proteomics to identify regulatory strategies and enzymatic homeostasis adaptations under nutrient limiting conditions (24, 25). Additional studies showed that *K. pneumoniae* responds to nutrient metal conditions via phosphorylation-based regulation (26, 27) and changes in kinase abundance (24, 25), confirming cellular modulation upon nutrient fluctuation. Beyond these areas, the intersection of nutrient acquisition, mass spectrometry-based proteomics, and microbial pathogenesis defines regulation of virulence factors in the human fungal pathogen, *Cryptococcus neoformans*, emphasizing the universality of these mechanisms to drive infection and disease (28, 29).

To expand upon these connections between nutrient acquisition and cellular regulation for *K. pneumoniae*, we performed mass spectrometry-based phosphoproteome profiling to identify and prioritize iron acquisition-associated modifications and regulatory proteins involved in promoting homeostasis. We defined iron-regulated phosphorylation of SbmA, a conserved homodimeric inner membrane protein associated with internalization of antimicrobial peptides and substrates (e.g., microcin B17, Bac7, bleomycin, microcin 25, peptide phosphorodiamidate morpholino oligomers and peptide nucleic acids (30–37)). Genetic deletion of *sbmA* in *K. pneumoniae* resulted in reduced cellular iron content and altered cell morphology, suggesting a role in iron acquisition. These putative functions were corroborated by proteome profiling of *ΔsbmA*, which showed compensatory induction of siderophore receptors and TonB-associated transport proteins, further supporting a role for SbmA in iron homeostasis. Integration with high-throughput drug screening identified a compound exploiting SbmA to inhibit bacterial growth, aligning with the promise of iron transport systems as effective antimicrobial targets(38). Collectively, our results demonstrate the combined power of phosphoproteomics and high-throughput screening for the discovery of novel antimicrobial agents that target critical bacterial pathways and uncovers a previously unrecognized role for SbmA in iron homeostasis.

## Material and Methods

### Bacterial strains and media preparation

Wild type (WT) *K. pneumoniae subsp. pneumoniae* (Kp52.145) was maintained on lysogeny broth (LB) agar plates and Δ*sbmA* was maintained on LB with 34 µg/mL chloramphenicol. M9 minimal media (LM) was made using DifcoTM M9 salts (6.78 g/L Na_2_HPO_4_, 3 g/L KH_2_PO_4_, 0.5 g/L NaCl, 1 g/L NH_4_Cl). Glucose was added to 0.4% (w/v), MgSO_4_ to 2 mM and CaCl_2_ to 0.1 mM. All glassware was rinsed with 3 M HCl to remove traces of iron or zinc, and solutions were made using Chelex® 100-treated (Bio-Rad) MilliQ water. Low salt Luria-Bertani LB media was made using 5 g/L yeast extract, 10 g/L tryptone, 0.5 g/L NaCl with 0.5 µM sterile EDTA. Preparation of strains, cultures, and *in vitro* experimentation performed as described (39), with study-specific modifications reported.

### Sample preparation for LC-MS/MS analysis

Four biological replicates of overnight cultures were established in 5 mL LB media in quadruplicate and incubated at 37 °C. Cells were pelleted by centrifugation at 3,500 x g for 10 min, then washed twice with 1 mL LM to remove any trace of LB media. Pelleted cells were reconstituted in 1 mL LM then subcultured 1:100 in 100 mL LM, LM supplemented with 10 µM iron (Fe_2_(SO_4_)_3_; iron-replete) and LM supplemented with 10 µM zinc (ZnSO_4_; zinc-replete) in quadruplicate. The cultures were incubated for 24 h at 37 °C (late stationary phase) then collected by centrifugation.

Pelleted cells were washed with phosphate-buffered saline (PBS) and then processed as previously described (40, 41), with some noted deviations. Briefly, cells were resuspended in 100 mM Tris-HCl pH 8.5, sodium dodecyl sulphate (SDS; 2%), proteinase inhibitor (Roche) and phosSTOP tablets (Roche). Cells were lysed using probe sonication (30 s on/ 30 s off in an ice bath, 30% power), reduced with 10 mM dithiothreitol and alkylated using 55 mM iodoacetamide. Proteins were precipitated using chloroform-methanol precipitation. Phase separation was achieved by adding 0.6 mL MilliQ water, then the top layer of the phase separation was removed carefully to avoid disrupting the white interphase containing protein. Methanol was added followed by vortexing and centrifuging at 10,000 x g for 5 min to pellet the protein. The resulting pellet was washed once with 0.4 mL methanol then dried in a SpeedVac at 45 °C for 10 min.

Protein pellets were resuspended in 8 M urea/ 40 mM HEPES buffer and the protein concentration was determined using BSA-tryptophan assay (42). Samples were digested overnight at room temperature using a 1:50 ratio of trypsin/lysC to protein then desalted using peptide desalting columns (Thermo Fischer Scientific, 89852), according to manufacturer’s instructions. The desalted peptides were split, with 10% (approx. 100 µg) dried to completion in SpeedVac at 45 °C and stored at -80 °C (total proteome samples). The remaining 90% (900 µg) were subjected to phosphopeptide enrichment.

### Phosphopeptide enrichment

Phosphopeptide enrichment was performed as previously described (40, 41, 43). Briefly, samples were reconstituted in 0.2 mL Binding/Wash buffer (Thermo Fisher Scientific, cat no A32992) and pH was adjusted to 2.5. Ferric nitrilotriacetate (Fe-NTA) column was equilibrated by washing twice with 0.2 mL Binding/Wash buffer then samples were incubated with a Fe-NTA column for 15 min (reduced from 30 min). The column was washed three times with 0.2 mL Binding/Wash buffer, once with 0.2 mL with MilliQ water then phosphopeptides were eluted using 0.2 mL Elution buffer. Phosphopeptides were dried completely in a SpeedVac at 45 °C.

### LC/MS-MS

Total proteome and phosphoenriched samples were reconstituted in 40 µL 0.1% formic acid (FA) and loaded onto Evotips according to the manufacturer’s instructions (44). Briefly, Evotip filters were wetted in 2-propanol, washed with 0.1 mL Buffer B (0.1% FA in ACN) (Fisher Scientific) then returned to 2-propanol and washed with 40 µL Buffer A (0.1% FA in water). Buffer A wash was repeated once, without returning tips to 2-Propanol. For the total proteome, 5 µg of protein was loaded per tip; for phosphoenriched samples, 90% of the sample volume was loaded per tip (36 µL). Samples were analyzed using a Themo Scientific Orbitrap Exploris 240 coupled with Evosep One liquid chromatography system. A 44 min method was used for phosphoenriched samples, and 88 min method was used for total proteome analysis. Mass spectrometry analysis was performed in data-dependent, positive ion mode using a 15 cm PepSep column with a 150 µm diameter and 1.9 µM beads (Evosep, EV1106). The precursor range was set at 350-1800 *m/z* with a resolution of 60,000, 50 msec injection time and an automatic gain control (AGC) target of 3e6. Dynamic exclusion was set to 10 sec, and charge states 2-8 were included. The fragment ion isolation window was 1 *m/z* with higher-energy collisional dissociation (HCD) fragmentation energy of 34 eV, a resolution of 15,000, and dynamic injection time. The first mass was set at 110 *m/z* for MS/MS scans, and the AGC target was set to 2e6.

### Raw data processing

The .RAW files were processed using MaxQuant v2.4.0.0 (45) with default parameters except where noted. Peptide identification was performed using the built-in Andromeda (46) search engine against the K52 *K. pneumoniae subsp. pneumoniae* proteome (5126 protein sequences from UP000000265 proteome ID; accessed Dec. 2, 2022). Phosphorylation was included as a variable modification using a modified version of the default phospho STY parameters, which included the addition of pHis and pAsp modifications (neutral loss of H_3_O_4_P; mass 97.9768950 Da) and a diagnostic peak for pHis (C_5_H_8_O_3_N_3_P immonium ion with a mass of 190.037604 *m/z* after protonation). Both modified and unmodified peptides were included for protein quantification using label-free quantification (LFQ) with the ratio count set to 1 (47). Other modifications included carbamidomethylation of cysteine as a fixed modification, methionine oxidation and N-acetylation of proteins as variable modifications. Digestion parameters were set to trypsin specificity with a maximum of 2 missed cleavages. The minimum number of peptides required to identify a protein was set to 2, and the “match between runs” feature was enabled with a match window time of 0.7 min and alignment time window of 20 min (47). Phosphopeptide abundance was normalized to the total proteome.

### Bioinformatics

Identification of phospho (STYDH) modifications and data visualization were performed in Perseus v2.0.10.0 (48) and ProteoPlotter (49). Phosphorylation multiplicities of 1, 2, and 3 were included. Phosphopeptides that matched contaminants, or the reverse database were removed, and the remaining phosphopeptides were classified into three groups: class I (localization probability >75%), class II (>50%) and class III (<50%). LFQ intensities were converted to the logarithmic scale (log_2_) then samples were filtered to exclude peptides identified in fewer than 3 replicates (3 valid values in at least one group).

### Generating competent K. pneumoniae cells

Competent *K. pneumoniae* cells were generated as previously described (50), with some noted deviations. Individual colonies of WT *K. pneumoniae* with pSim6 plasmid were used to inoculate 5 mL LB media (with 100 µg/mL ampicillin) overnight at 30 °C. The next day, 0.5 mL of culture was used to inoculate 50 mL of low salt LB media which was incubated at 30 °C for 5.5 hr until OD_600nm_ was between 0.4 - 0.5. Next, the pSim6 promoter was activated at 42 °C for 1 h, cells were washed three times with ice-cold sterile MilliQ water then washed with 1 mL sterile, ice-cold 10% glycerol and reconstituted with 0.4 mL ice-cold, sterile 10% glycerol.

### Deletion Strain Generation

Deletion of *sbmA* was performed using the lambda-red recombinase system (51), as we previously described (24). First, the chloramphenicol resistance gene was amplified from the pkD3 plasmid with overhangs complementary to the first and last 50 nucleotides of *sbmA*. The PCR product was added to 50 µL competent *K. pneumoniae* then electroporated at 1800 V. Electroporated cells recovered in 0.5 mL LB media at 37 °C for 3 h followed by overnight incubation at room temperature. Recovered cells were plated on half-strength chloramphenicol LB plates (17 µg/mL chloramphenicol) at 50 µL, 150 µL, and 300 µL and incubated overnight at 37°C then colony PCR was performed to identify deletion strains. PCR primer design is provided (Table S1).

### PCR Screening of deletion strain colonies

Colonies were screened using size-shift colony PCR to confirm gene knockout. Primers recognizing 100 nucleotides upstream/downstream of the gene of interest were used, and positive deletion strains were identified by a shift in gene size from *sbmA* gene size (1221 bp) to chloramphenicol (1031 bp). PCR primer design is provided (Table S1). Positive colonies from size shift PCR were screened using inside-outside colony PCR to confirm the presence and location of chloramphenicol resistance gene insert. WT was used as the negative control. PCR primer design is provided (Table S1). Whole genome sequencing was performed to confirm *sbmA* deletion (Table S2).

### Protein structure prediction

The AlphaFold2 predicted structure of Kp-SbmA (A0A2S8TRL6) was used as a query in the DALI server (http://ekhidna2.biocenter.helsinki.fi/dali/) to identify structurally similar proteins with experimentally resolved crystal structures. Based on the oligomeric states observed in these homologs, two additional structural models were generated using AlphaFold3 (52). One model represented the native (non-modified) state, and the other included a phosphorylation modification at residue Y285. To determine spatial arrangement within the inner membrane of *K. pneumoniae*, the models were analyzed using the Orientations of Proteins in Membranes (OPM) database (https://opm.phar.umich.edu). Final visualizations and structural superimpositions were performed using PyMOL v3.1.5.1.

### Growth curves

To assess differences in growth, overnight cultures of WT and Δ*sbmA* were subcultured 1:100 into 100 µL LB, LM, LM + 10 µM Fe_2_(SO_4_)_3_ and LM + 10 µM ZnSO_4_ media in clear, sterile 96-well plates. Cultures were grown for 24 h at 37 °C with shaking and OD_600 nm_ was measured every 15 min using a BioTek plate reader. Data were collected in biological triplicate and technical duplicate and plotted using GraphPad Prism v10.

### Cellular iron quantification assay

Overnight cultures of WT and Δ*sbmA* were subcultured 1:100 into minimal media and allowed to grow to mid-log (5 h) at 37 °C in biological quadruplicate. Then 10^9^ cells were collected and washed twice with 0.4 mL 0.9% NaCl followed by reconstitution in 0.2 mL lysis buffer. Iron ions are reduced to ferrous state. After cells were lysed, 80 µL supernatant was collected and transferred to a clear 96-well plate with 80 µL chromogenic solution and incubated at 37 °C for 40 min. Absorbance was measured using a Biotek plate reader at 593 nm and total iron was calculated using an iron standard curve (Fig. S1). Data was plotted and analyzed using GraphPad Prism v10.

### Macrophage infection

Immortalized BALB/c macrophages (generously provided by Dr. Felix Meissner, from University Hospital Bonn) were infected with *K. pneumoniae* strains as previously described (53, 54) with some noted deviations. Macrophage cells were seeded on 24-well plates at 5 x 10^4^ cells per well in Dulbecco’s modified Eagle’s medium (DMEM) (Glutamax DMEM, 10% FBS, 2 mM Glutamax, 1% sodium pyruvate, 1% L-glutamine, 5% pen/strep) for 48 h at 37 °C and 5% CO_2_. After 48 h, macrophages achieved confluence (∼80% well coverage; 5 x 10^5^ cells/mL), DMEM media was removed, cells were washed twice with 1 mL room temperature, sterile phosphate-buffered saline (PBS), then WT and Δ*sbmA* were added at a multiplicity of infection (MOI) of 100:1 (5 x 10^7^ bacteria cells/mL in DMEM without antibiotics) in triplicate. Three replicates of uninfected macrophages were also grown in 1 mL DMEM without antibiotics. Cells were incubated at 37 °C and 5% CO_2_ for 90 min.

### Bacterial cell counting

Cell counts were performed for each strain at mid-log (3 h for WT and 2 h for Δ*sbmA*) using a hemocytometer. Standard curves for WT (Fig. S2a) and Δ*sbmA* (Fig. S2b) were generated by dilution series from 2X to 256X, using sterile PBS in a sterile 96-well plate. OD_600nm_ was measured for each dilution series and standard curves were fitted with a linear trendline. The remaining replicates were diluted 10X, OD_600nm_ was measured, and cell number per mL was calculated using the corresponding standard curve equation for each strain.

### Lactate dehydrogenase cytotoxicity assay

Following co-culture of *K. pneumoniae* and macrophages, cytotoxicity was measured using Promega CytoTox 96® Non-Radioactive Cytotoxicity Assay (Promega, G1780) according to manufacturer instructions. Uninfected samples served as controls with quantification performed at OD_490nm_. Data were plotted and analyzed using GraphPad Prism v10.

### Phagocytosis evasion and enumeration of phagocytosed bacterial

Following co-culture, macrophages were burst using 1.2% triton to release internalized bacteria. This suspension was diluted 10X from 10^0^ – 10^4^ with 100 µL of each dilution plated on LB agar and incubated overnight at 37 °C. Individual colonies were counted to determine colony forming units (CFU). Data were plotted and analyzed using GraphPad Prism v10.

### Transmission electron microscopy

Overnight cultures of WT and Δ*sbmA* cultures were collected at 4000 rpm for 5 min then washed twice with 5 mL 50 mM HEPES buffer and fixed for 1 h in 2.5% glutaraldehyde 2% paraformaldehyde. Cells were washed twice with 50 mM HEPES then enrobed in noble agar (final concentration 2%) and spread as a thin sheet to generate 1 mm thick agar slices. Agar slices were then treated with 1% osmium tetroxide for 45 min to increase sample contrast. Samples were dehydrated using an EtOH dehydration series (25%, 50%, 75%, 95%, 100%), incubated in LR white Resin for 1 h then transferred to capsules and polymerized in LR white overnight at 60 °C. Polymerized capsules were then prepared for sectioning using Reichert-Jung Ultracut E Ultramicrotome then 88 nm sections were generating using a diamond blade. Sections were collected on formvar coated grids then stained for 5 min in 5% uranyl acetate then stained in lead citrate for 10 min. Samples were then imaged using 120kV field emission transmission electron microscope (FEI Tecnai G2 F20).

### Scanning electron microscopy

Overnight cultures of WT and Δ*sbmA* were collected at 4000 rpm for 5 min then washed twice with 1 mL phosphate buffer. Samples were resuspended in 0.4 mL phosphate buffer with half the sample added adhered onto carbon planchet for 30 min followed by 2% glutaraldehyde fixing for 30 min. Samples were washed three times with phosphate buffer and submerged in 1% osmium tetroxide for 30 min to increase sample contrast. Samples were dehydrated using an EtOH dehydration series (50%, 70%, 80%, 90% then 100% ethanol). Critical point drying (Denton DCP-1) was used to replace EtOH with CO_2_ then sputter-coating with gold and palladium (Denton Desk V TSC; 90 s under argon atmosphere). Images were captured using scanning electron microscopy (FEI Quanta FEG 250).

### Biofilm crystal violet assay

Overnight cultures of WT and Δ*sbmA* were subcultured 1:100 into nutrient-rich tryptic soy broth (TSB) media until mid-log (3 h and 2 h, respectively). Cells were counted and cultures were diluted to 10^8^ cells/mL followed by 1/10 diluted into 200 µL TSB or LM media per well in a clear, sterile 96 well plate. Diluted cultures were washed twice with 1 mL LM before subculturing into LM media. Plates were incubated statically for 30 h to achieve mature biofilms. Biofilms were washed twice with 200 µL PBS, allowed to dry, then 200 µL 0.2% crystal violet was added to each well for 10 min, washed twice with 200 µL water, and allowed to dry. Crystal violet was resuspended with 200 µL 30% acetic acid for 10 min then 100 µL was transferred to a new 96 well plate. Absorbance was measured at OD_550nm_ to quantify biofilm production. Biofilm assays were performed in biological triplicate and in technical duplicate.

### Biofilm resuspension assay

Resuspension assays were set up as described for crystal violet assay, except in place of crystal violet dye, 200 µL PBS was added and plates were sonicated for 2 min to resuspend biofilms. Resuspended biofilms were diluted by factor 10 from 10^4^ - 10^7^ for TSB cultured biofilms and from 10^2^ – 10^5^ for LM cultured biofilms. Next, 100 µL of diluted resuspended biofilm was added to an LB plate and allowed to grow for 24 h. The next day, individual colonies were counted and CFU counts were determined. The experiment was performed in biological triplicate and technical duplicate.

### Compound library screening

A high throughput screen of a small-molecule compound library was conducted on WT and Δ*sbmA* strains at SPARC Drug Discovery at The Hospital for Sick Children. The library contained 2,500 drug-like compounds exhibiting extensive pharmacophore coverage and chemical diversity. Overnight cultures were diluted 1:100 in LB media, then 50 µL of these subcultures were dispensed into each well of 384-well plates containing pre-dispensed compounds (dissolved in DMSO) to obtain a final concentration of 10 µM. The plates were incubated for 24 h at 37 °C with shaking at 200 rpm. OD_600nm_ values were measured using a microplate reader, and the average growth and standard deviations were calculated for sample and control populations. Hit confirmation was performed by repeating the assay in triplicate for compounds that reduced culture growth by more than 2 standard deviations from the negative control mean in the primary screen. Kanamycin (50 µg/mL) wells were included as positive controls and DMSO as negative vehicle controls. Data were archived and analyzed using the CDD Vault database from Collaborative Drug Discovery (Burlingame, CA. www.collaborativedrug.com).

### Compound growth inhibition assay

WT and Δ*sbmA* were cultured overnight then diluted 1:100 into Mueller Hinton (MH) broth, the standard growth media for antibiotic susceptibility assays. Next, 1-(3-bromophenyl)-3-(4-fluorophenyl)urea (compound 422), was added at 20 µM – 0.625 µM, serially diluted, in a sterile polystyrene 96-well plate and incubated with shaking for 24 h at 37 °C. WT and Δ*sbmA* with no compound were included to determine maximum growth and uninfected MH media was included to determine minimum growth. OD_600nm_ was measured at 24 h and growth was normalized where 100% was the OD_600nm_ value when no compound was added and 0% was the uninfected MH media OD_600nm_ value. This assay was performed in biological quadruplicate and technical duplicate.

### Compound cytotoxicity and CFU counting in macrophage

As outlined above, macrophage were incubated in the presence of compound 422 at 10 and 20 µM, along with an untreated control followed by quantification of LDH release to assess potential cytotoxicity. CFU counts from supernatant and phagocytosed cells were performed as described above for a 90 min-co-culture in the presence of compound 422 at 1, 10, or 20 µM.

### Statistics and reproducibility

Comparisons between two or three data sets were performed using unpaired two-tailed Student’s *t*-tests and comparisons between four or more data sets were performed using a one-way ANOVA. Significance annotations were applied as follows: *p < 0.0332, **p < 0.0021, ***p < 0.0002 and ****p < 0.0001. For proteomic analysis missing values were imputed using a Gaussian distribution with a standard deviation width of 0.3 and a downshift of 1.8. A Student’s *t*-test was conducted to identify significant changes in phosphoprotein abundance in different conditions using a *p* ≤ 0.05, a Benjamini–Hochberg false discovery rate (FDR) of 5% and an S_0_ value of 1. Volcano plot values were normalized by subtracting the median from columns to center data at zero.

## Results

### Iron availability influences phosphorylation of transport and biosynthesis proteins in K. pneumoniae

Metal acquisition is essential for supporting a wide array of cellular functions (10, 55, 56) and phosphorylation serves as a key regulatory mechanism for these processes (57). Despite their importance, the intersection of metal availability and phosphorylation remains underexplored in *K. pneumoniae*. To address this gap, we systematically identified proteins whose phosphorylation status is modulated by iron and zinc availability with the goal of uncovering regulatory nodes in metal homeostasis that could serve as novel targets for antimicrobial intervention. In total, we identified 640 phosphopeptides, of which 574 were class I (>75% localization probability) mapping to 315 phosphoproteins that were associated with diverse biological processes in a media-dependent manner. Under iron-replete conditions (Fig. 1a), proteins involved in biosynthetic processes became more abundantly phosphorylated (22% of the phosphoproteins identified) compared to the iron-limited conditions (12% for zinc-replete and 13% for LM; Fig. 1b-c). We observed that proteins involved in transport were two-to-three-fold more abundantly phosphorylated under iron-limited conditions (i.e., 10% of phosphorylated proteins in zinc-replete media [Fig. 1b], 7% in LM [Fig. 1c], and only 3% in iron-replete media [Fig. 1a]).

**Figure 1:**
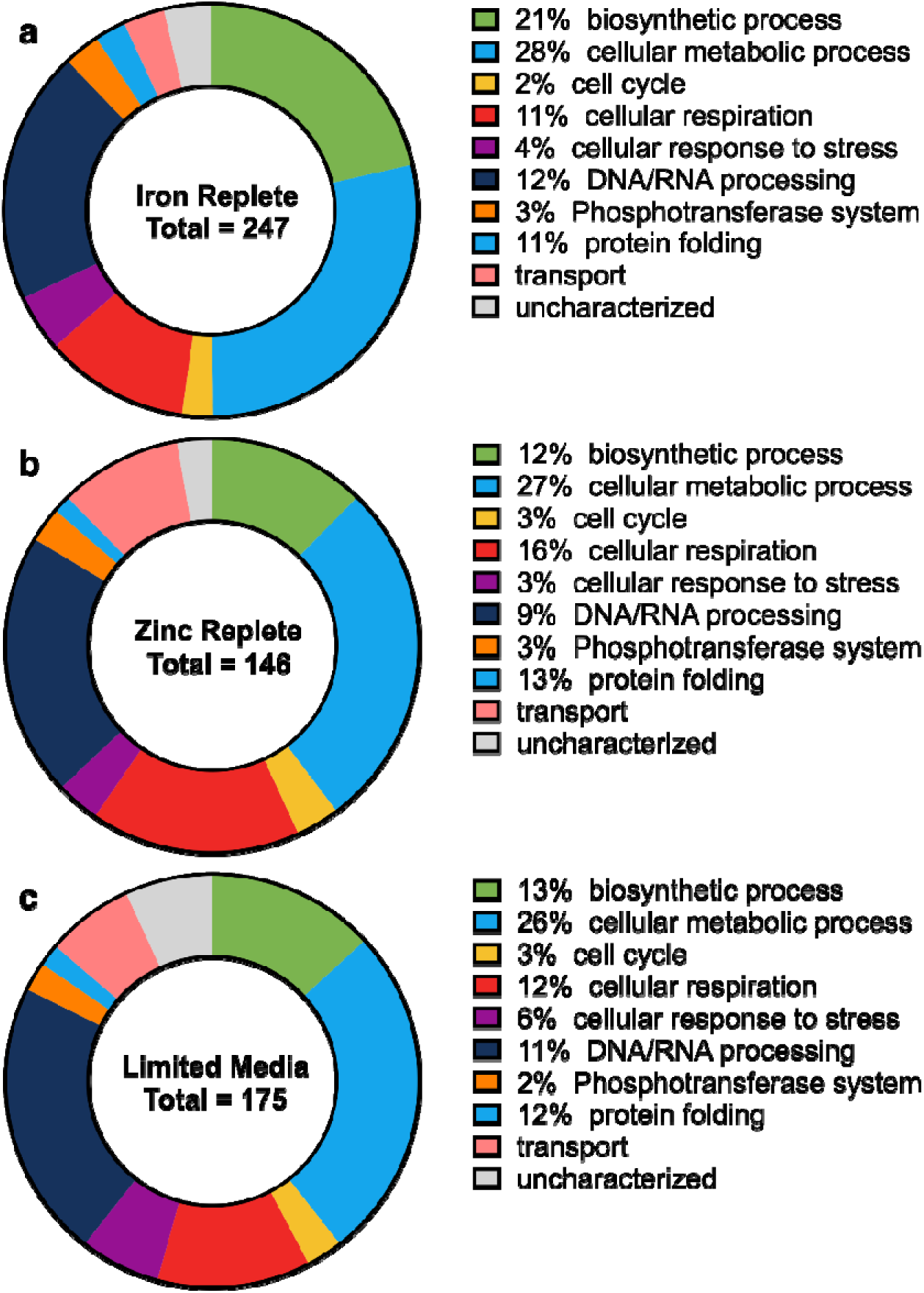
Distribution of phosphoproteins from mass spectrometry profiling of *K. pneumoniae* under modulated media conditions. Phosphoproteins organized by Gene Ontology Biological Processes under **(a)** iron-replete; **(b)** zinc-replete; and **(c)** limited media (LM) conditions. Experiment performed in biological quadruplicate. Class I phosphosites (localization probability >75%).

### Prioritization of transport-associated proteins identified SmbA with putative roles in metal acquisition and confirmed phosphorylation

To investigate potential links between phosphoregulation and iron transport, we highlighted 15 transport proteins that were more abundantly phosphorylated under iron-limited conditions (i.e., LM or zinc-replete condition) (Table 1). As expected, many prioritized proteins have established roles in the transport of amino acids, proteins, and proteins (SecD (58), SecE (59), SecF (59), YliA (60), PrlA (61)) or other essential nutrients, such as carbohydrates (MglA (62)), sulfate (Sbp (63)), sodium and potassium (PutP (64), KPN_00790 (65)), iron sulfur cluster assembly (SufC), or cell membrane stability (OmpA, OmpC). Regulation of these systems under iron-limited conditions was expected, given iron’s role as a cofactor in protein and amino acid biosynthesis (66). Our focus, however, turned to SbmA, a transport protein involved in the internalization of microcins (67), which exploit nutrient transport systems via a “Trojan horse” strategy (68), but have no known role in iron acquisition and/or transport. Given prior research suggesting that iron uptake pathways can be similarly hijacked by naturally occurring antimicrobial compounds (69–71), we assessed SbmA protein interaction networks and observed multiple connections with proteins involved in siderophore biosynthesis and regulation (KPN_00318 (72), YpfG (65, 73), YaiW (74)) and iron uptake (KPN_00122 (75, 76)) (Fig. 2a). Notably, we performed a BLASTp homology search on K52 *K. pneumoniae* SbmA compared to SbmA homologues (maximum target species of 5,000), resulting in classification across nine genera based on taxonomic rank with average shared sequence identity at 89%, supporting strong functional or evolutionary constraint of SbmA (Fig. S4).

**Figure 2:**
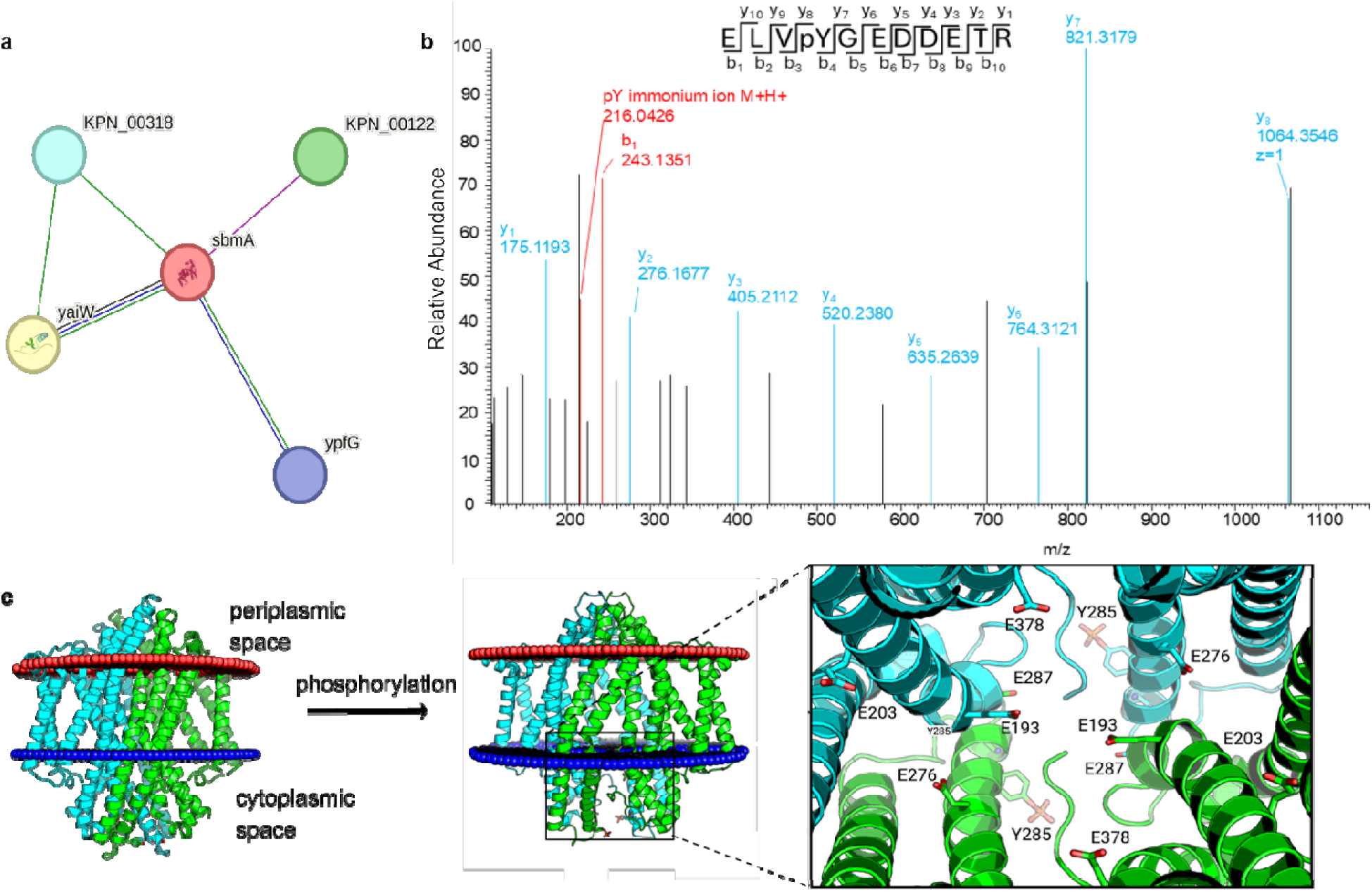
Computational analysis of SbmA network, phosphosite localization, and predicted structure. **(a)** Predicted protein-protein interaction network from STRING database. Known interactions are indicated by fuchsia (experimentally determined) or turquoise (from curated databases) and predicted interactions are indicated by green (gene neighbourhood), blue (gene co-occurrence) and purple (co-expression). **(b)** HCD spectra of *K. pneumoniae* SbmA phosphopeptide found in zinc-replete conditions for ELVpYGEDDETR peptide corresponding to Y285 phosphosite. **(c)** AlphaFold structure prediction with highlighted Y285 phosphosite and glutamate ladder, designed in PyMOL.

**Table 1:** Phosphorylated transport proteins identified in LM and/or zinc-replete media with associated functions (gathered from UniProt(65), based on GO annotations)

| **Gene** | **Media** | **Function** |
| --- | --- | --- |
| *sbmA* | Zinc | Microcin B17 transport |
| *secF* | Zinc | protein transport |
| *secE* | Zinc | protein transport |
| *tauC* | LM & Zinc | Taurine transport |
| *secD* | LM & Zinc | protein transport |
| KPN_00790 | LM & Zinc | sodium/potassium transport |
| *yliA* | LM | Glutathione import |
| *ompA* | LM & Zinc | cell membrane stability |
| *sufC* | LM | ATP-binding component of a transport system |
| *mglA* | LM & Zinc | carbohydrate import |
| *yojI* | LM | peptide transport |
| *ompC* | LM & Zinc | porin |
| *prlA* | LM & Zinc | protein transport |
| *sbp* | LM & Zinc | Sulfate transport |
| *putP* | LM & Zinc | Sodium/proline symporter |

Next, we confirmed phosphorylation of SbmA under iron-limited conditions at the Y285 residue in the ELVpYGEDDETR peptide (Fig. 2b). Based on cryo-electron microscopy analysis of SbmA in *Escherichia coli* (67) (with a Root mean square deviation to the *K. pneumoniae* homologue of 1.054; Fig. S5), the Y285 residue is cytosolic-facing; however, phosphorylation of this residue is predicted to trigger a conformational change that opens the cytoplasmic-facing gate of the transport protein to potential enable substrate transport (Fig. 2c). The predicted template modelling score (pTM) of SbmA in the unphosphorylated form is 0.7 and 0.8 in the phosphorylated form, indicating a high confidence level in the AlphaFold predicted structures (77–79) and a putative effect of iron limitation on protein function. These findings reveal that iron-dependent phosphorylation of SbmA may influence its function as a regulatory node in iron acquisition.

### ΔsbmA strain influences bacterial intercellular iron content, growth, virulence, and phagocytosis in an iron-dependent manner

To investigate a connection among SbmA phosphorylation, media conditions, and bacterial survival, we generated a *sbmA* deletion strain and quantified cellular changes. To support the putative role of SbmA in iron acquisition as predicted by a confirmation change upon phosphorylation, we assessed if the protein has a role in iron acquisition by quantifying total cellular iron for WT and Δ*sbmA* strains cultured in LM (i.e., iron-limited media). This assay determined that Δ*sbmA* had an average of 176.4 pmol iron (Fe^2+^ and Fe^3+^) per 10^9^ cells compared to 579.8 pmol for WT, indicating three-fold less internalized iron in the Δ*sbmA* strain compared to WT (Fig. 3i; *p* = 0.0136) (Fig. 3a). This is the first reported connection between SbmA and cellular iron content and corroborates our phosphoproteome profiling and predictive functional properties.

**Figure 3:**
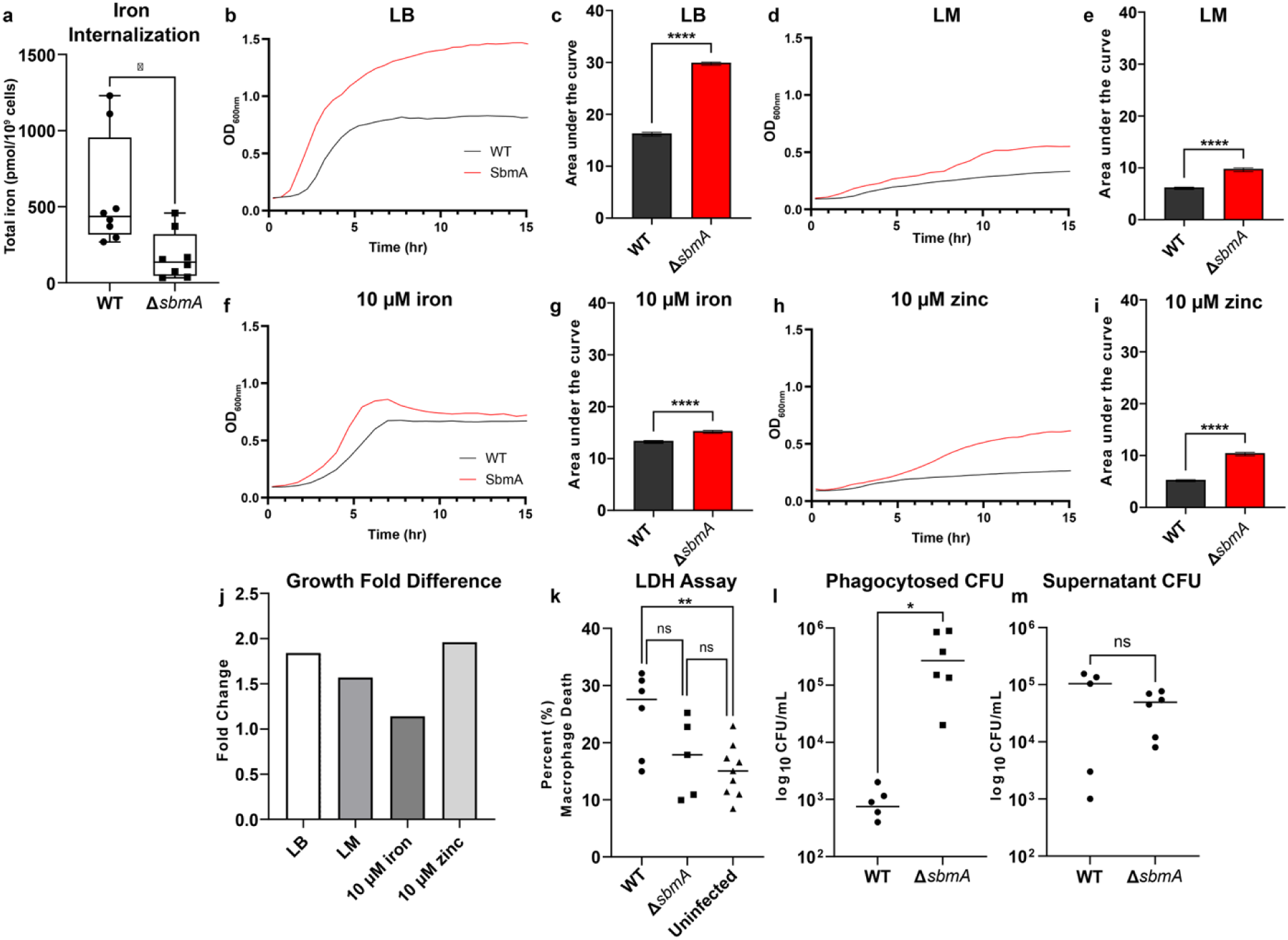
Phenotypic characterization by iron internalization, growth, and virulence of WT (black) and Δ*sbmA* (red). **(a)** Total iron content (Fe^3+^ and Fe^2+^) of strains grown in LM in biological quadruplicate and technical duplicate. Box and whisker plot illustrates the median, second and third quartile and minimum and maximum values of each data set. **(b)** 15 h growth curve in LB media. **(c)** Area under the curve (AUC) for LB growth curve. **(d)** 15 h growth curve in LM. **(e)** AUC for LM growth curve. **(f)** 15 h growth curve in LM + 10 µM Fe_2_(SO_4_)_3_. **(g)** AUC for LM + 10 µM Fe_2_(SO_4_)_3_ growth curve. **(h)** 15 h growth curve in LM + ZnSO_4_**. (i)** AUC for LM + ZnSO_4_ growth curve. **(j)** Fold-difference of growth across media conditions. Optical density (OD) was measured at 600 nm in biological triplicate and technical duplicate and curves are a representative of the mean. AUC bars indicate the mean and error bars denote standard error (N=6). **(k)** Lactate dehydrogenase (LDH) cytotoxicity assay for WT and Δ*sbmA* infected BALB/c macrophages and uninfected macrophages after 90 min co-culture in biological triplicate and technical duplicate. Data is displayed as percentage of max death. **(l)** Colony forming units (CFU) from WT and Δ*sbmA* phagocytosed by BALB/c macrophages during 90 min co-culture in biological triplicate and technical duplicate. (,) CFU for WT and Δ*sbmA* in the supernatant following BALB/c 90 min co-culture. LDH and CFU plots display mean as a line. Statistical significance determined using a two-tailed Student’s *t*-test where *p < 0.0332, **p < 0.0021, ****p < 0.0001.

Next, we assessed the impact of *sbmA* deletion on growth dynamics under multiple media conditions. Under nutrient-rich LB conditions, Δ*sbmA* entered exponential phase approximately one hour earlier than WT and reached a higher final optical density (OD_600nm_), resulting in an approximately two-fold increase in growth compared to WT (Fig. 3b, 3c; *p* < 0.0001). Under minimal media (LM) conditions, Δ*sbmA* exhibited a more modest growth enhancement, with a slightly elevated final OD_600nm_ corresponding to a 1.5-fold increase in growth compared to WT (Fig. 3d, 3e; *p* < 0.0001). Under iron-supplemented LM conditions, Δ*sbmA* reached exponential phase 0.5–1 hour earlier than WT and achieved a 1.1-fold higher growth compared to WT (Fig. 3f, 3g; *p* < 0.0001). Under zinc-supplemented conditions, Δ*sbmA* reached a higher final OD_600nm_, corresponding to a two-fold increase in growth compared to WT (Fig. 3h, 3i; *p* < 0.0001). These findings support dysregulated growth for Δ*sbmA* under altered media conditions with a reduced differential in the absence of iron (Fig. 3j).

Given our recent observation that altered iron acquisition impacts bacterial virulence and evasion of macrophage phagocytosis (80), we evaluated the effect of Δ*sbmA* on bacterial virulence within an *in vitro* model. For this assessment, we performed a macrophage infection assay using WT and Δ*sbmA* strains to determine host cell death via lactate dehydrogenase (LDH) release. We observed a 7.63% decrease in host cell death upon co-culture with Δ*sbmA* compared to WT, however this difference was not statistically significant (*p* = 0.1113; Fig. 3k). We hypothesized this decrease in virulence may be attributed to the decreased ability for Δ*sbmA* to obtain iron from its environment, an important virulence factor for bacteria (80, 81). Notably, cell death was significantly higher upon WT co-cultured versus uninfected macrophages (*p* = 0.0064) whereas there was no significant difference in cell death between Δ*sbmA* co-cultured and uninfected macrophages (*p* = 0.4656). We also assessed potential differences in bacterial phagocytosis between strains and observed a 483-fold increase in the number of phagocytosed Δ*sbmA* cells compared to WT (*p* = 0.0043; Fig. 3l), despite only a 2-fold reduction Δ*sbmA* cells remaining in the supernatant (*p* = 0.3009; Fig. 3m). These findings suggest that macrophages more readily internalize Δ*sbmA* (corresponding with our previous work linking iron acquisition to phagocytic evasion(80)) and yet, are less effective at clearing the pathogen once engulfed, supporting the exploitation of SbmA by macrophages to deliver antimicrobial compounds to clear *K. pneumoniae* infection.

### ΔsbmA alters bacterial morphology, promotes cell clumping, and increases biofilm production

Given the role of nutrients, such as iron, in cell morphology (82–84) we applied electron microscopy to visualize differences in cell size and shape between Δ*sbmA* and WT. Based on prior reports, we hypothesized that Δ*sbmA* would display a filamentous morphology due to iron-stress (82, 84) and display reduced cell size due to impaired iron-dependent biosynthetic activity. Using transmission electron microscopy (TEM; high-resolution images of individual cells), we observed a clear morphological distinction between strains with WT cells maintaining a rod-shaped morphology (Fig. 4a), while Δ*sbmA* cells appeared spherical (Fig. 4b), suggesting a stress-induced morphological adaptation. To validate these findings, we performed scanning electron microscopy (SEM; detailed visualization of surface structures and cell communities), and as expected, we observed spherical morphology of Δ*sbmA* cells, along with additional unique characteristics, including cell clumping and filamentation (Fig. 4c), in contrast to the dispersed single-cell morphology of WT (Fig. 4d). From the TEM images, we quantified these morphological differences by measuring cell length and width determining an average length 1.16 µm for Δ*sbmA* cells compared to 1.93 µm for WT, corresponding to a 40% reduction (*p* < 0.0001; Fig. 4e). Cell width was also reduced at 0.77 µm for Δ*sbmA* cells compared to 0. 0.98 µm for WT (*p* < 0.0001; Fig. 4f). We observed corroborating findings from SEM measurements with cell length of 1.42 µm for Δ*sbmA* cells compared to 2.48 µm for WT (*p* < 0.0001; Fig. 4g), and widths of 0.76 µm for Δ*sbmA* cells compared to 0.93 µm for WT (*p* < 0.0001; Fig. 4h).

**Figure 4:**
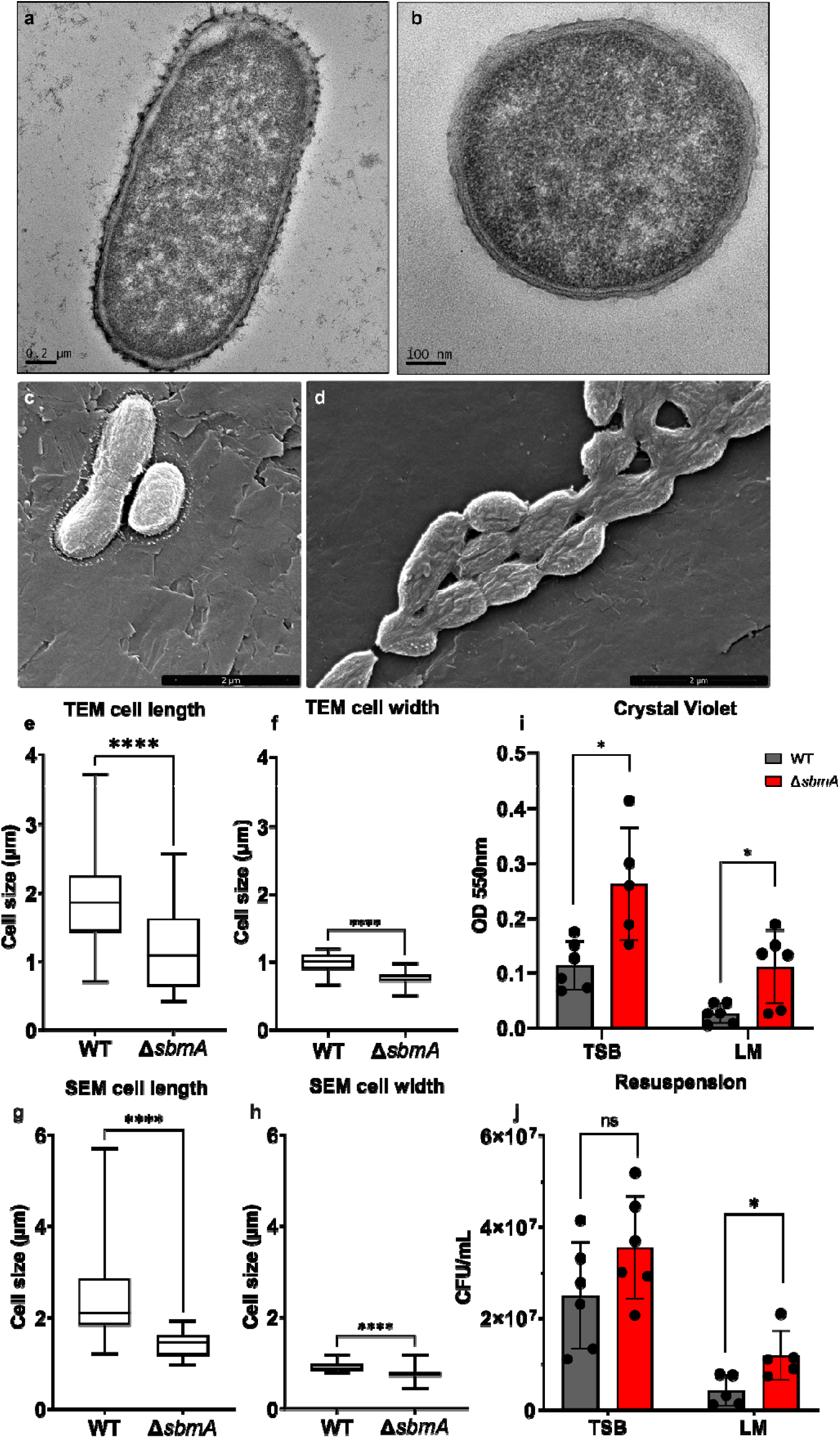
Cell morphology visualization by TEM and SEM and biofilm quantification. **(a)** Transmission electron microscopy (TEM) WT micrograph. **(b)** TEM Δ*sbmA* micrograph. **(c)** Scanning electron microscopy (SEM) 50,000X magnification WT micrograph. **(d)** 50,000X magnification Δ*sbmA* micrograph. **(e)** TEM cell length for WT (N=45) and Δ*sbmA* cells (N=43). **(f)** TEM cell width for WT (N=47) and Δ*sbmA* (N=47). **(g)** SEM cell length for WT (N=47) and Δ*sbmA* (N=64). **(h)** SEM cell width for WT (N=30) and Δ*sbmA* (N=30). Strains were cultured in LB media and collected at stationary phase for TEM and mid-log for SEM. Images are representative of the general morphology from ≥20 captured images from ≥5 fields of view. Box and whisker plots show the median, second and third quartile and minimum and maximum values. **(i)** Crystal violet (CV) biofilm staining and quantification at OD_550 nm_ after 30 h static incubation. Performed in biological triplicate, with 7 replicates (averaged), in technical duplicate. **(j)** Biofilm resuspension and CFU quantification. Performed in biological triplicate and technical duplicate. Error bars denote standard deviation. Statistical differences between strains were calculated using a two-tailed Student’s *t*-test where *p < 0.032 and ****p < 0.0001.

Next, we hypothesized that cell clumping, which increases surface area, of *ΔsbmA* cells would promote biofilm formation, given the critical role of surface binding in this process(85). To evaluate an impact of cell morphology and nutrient availability on biofilm formation, we cultured Δ*sbmA* and WT strains in nutrient-rich (TSB) or nutrient-limited (LM) media for 30 h to allow for mature biofilm development (86). Crystal violet (CV) staining, which quantifies total biofilm biomass, revealed that Δ*sbmA* produced significantly more biofilm than WT, with a two-fold increase in TSB conditions (*p* < 0.0001; Fig. 4i) and a four-fold increase in LM conditions (*p* = 0.0057; Fig. 4i). These results showed similar trends by the resuspension assay, with a 1.5-fold increase for Δ*sbmA* in TSB conditions (*p* = 0.1434) and a three-fold increase for Δ*sbmA* in LM conditions (*p* = 0.0261) compared to WT (Fig. 4j). The reductions in both cell length and width are consistent with nutrient stress and diminished biosynthetic capacity under iron-limited conditions (87). Further, the clumping phenotype observed in Δ*sbmA,* which also correlated with biofilm production, is consistent with iron-stress responses previously described in Gram-negative bacteria (82, 84). These findings support a model in which SbmA contributes to iron acquisition, and gene deletion triggers compensatory morphological changes, such as filamentation and cell clumping, resulting in increased surface area and biofilm adhesion.

### Proteome profiling confirms the role of SbmA in metal acquisition and compensatory functional activation

To further validate the role of SbmA in iron acquisition, we performed proteome profiling of Δ*sbmA* and WT cultured in LM media at mid-log phase. Across all samples, we identified 1,974 cellular proteins, with a principal component analysis (PCA) revealing distinct clustering by strain (component 1; 52%) and biological replicate (component 2; 19.6%) (Fig. 5a). Differential abundance analysis identified 244 proteins with significantly higher abundance in WT and 369 proteins with significantly higher abundance in Δ*sbmA* (Fig. 5b). GOBP annotation mapping of the significantly different proteins, revealed that iron-associated proteins were approximately two-fold less abundant in Δ*sbmA* compared to WT, whereas other metal cofactors, including Mg, Mn, Cu, and Zn, were more prevalent in Δ*sbmA* compared to WT, supporting compensatory regulation upon reduced iron acquisition properties (Fig. 5c). As a percentage of significant different proteins, these trends were consistent: iron co-factor proteins accounted for 5.3% of significant higher proteins in WT compared to 1.9% in Δ*sbmA*, and other co-factors accounted for 9.0% in WT compared to 11.4% in Δ*sbmA* (Fig. 5d, 5e).

**Figure 5:**
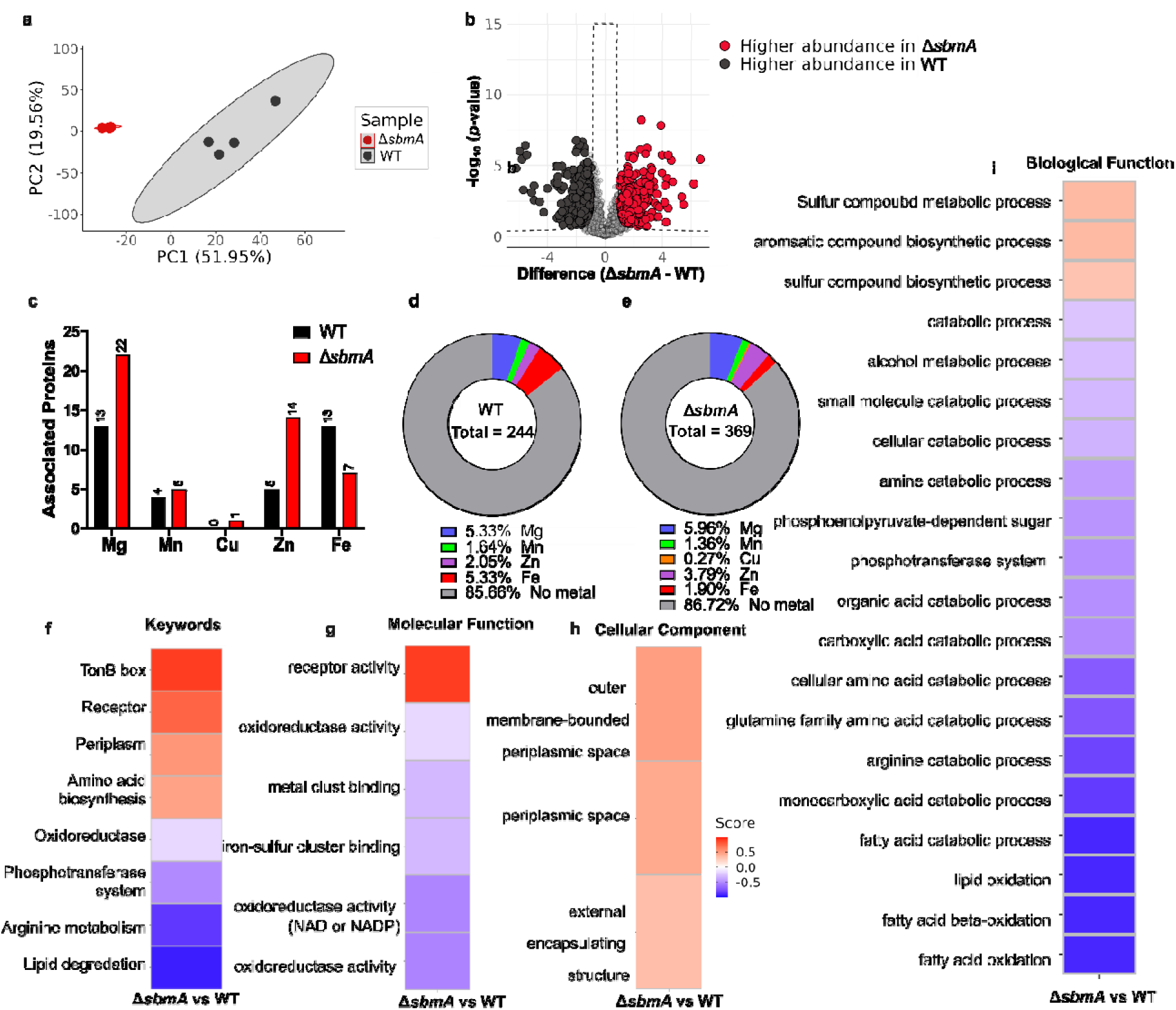
Proteome profiling on WT and Δ*sbmA* cultured to mid-log in limited media (LM). **(a)** Principal component analysis of sample variability between Δ*sbmA* and WT. **(b)** Volcano plot depicting all proteins identified in Δ*sbmA* and WT. Data was normalized by subtracting median from columns. Significantly higher abundance in Δ*sbmA* = red; significantly higher abundance in WT = black. **(c)** Number of proteins significantly more abundant in WT vs Δ*sbmA* with known metal cofactors in LM by GO annotations. **(d)** Percentage of proteins with known metal cofactors that are significantly more abundant in WT. **(e)** Percentage of proteins with known metal cofactors that are significantly more abundant in Δ*sbmA*. **(f)** One-Dimensional (1D) annotation keyword enrichment of proteins from Δ*sbmA* vs WT. **(g)** 1D annotation for GOMF enrichment. **(h)** 1D annotation for GOCC enrichment. **(i)** 1D annotation for GOBP enrichment. Scores approaching 0.5 (red) indicate that the term is over represented in Δ*sbmA* whereas scores approaching -0.5 (blue) indicate that the term is under represented in Δ*sbmA*. Significance was determined using a Student’s *t*-test with *p*-value ≤ 0.05. False discovery rate (FDR) = 0.05 and S_0_ = 1. Generated using ProteoPlotter.

To further investigate the functional consequences of *sbmA* deletion, we performed 1D annotation enrichment analysis using Keywords and GO terms (GOBP, GOMF, GOCC). Proteins associated with receptor activity, periplasmic space, and the outer membrane were significantly more abundant in Δ*sbmA* than WT (Fig. 5f–h). Notably, receptor proteins enriched in Δ*sbmA* were almost exclusively siderophore receptors, including *fhuA* (3-fold), *fepA* (4-fold), *yncD* (5-fold), KPN_02304 (7-fold), and KPN_01108 (15-fold), which mediate the import of ferrichrome, enterobactin, ferrioxamine, and aerobactin (Table S3) (65, 88, 89). Δ*sbmA* also showed increased abundance of TonB box–containing proteins (Fig. 5f), which facilitate iron transport across the outer membrane (90). Conversely, WT cells were enriched in proteins involved in metal and iron-sulfur cluster binding, correlating with higher intracellular iron availability, oxidoreductase activity for redox homeostasis, and core metabolic processes, such as alcohol, amine, and lipid metabolism, all of which utilize iron as a cofactor (91–93) (Fig. 5g, 5i). The enrichment of iron-associated proteins in WT compared to compensatory enrichment observed for Δ*sbmA* cells supports our theory that cells adapt to the loss of SbmA by using alternative metal acquisition pathways, reinforcing the link between the protein and iron homeostasis.

### High-throughput drug screening identifies a compound that exploits SbmA to reduce bacterial growth with minimal host cytotoxicity

Given the role of SbmA in antimicrobial internalization (67), a function leveraged by macrophage for pathogen destruction, we used high throughput drug screening to identify compounds that exploit SbmA to inhibit *K. pneumoniae* growth. To achieve this, we evaluated a 2,500 drug-like compound library for reduced growth of WT, but not Δ*sbmA*. Our screening identified 1-(3-bromophenyl)-3-(4-fluorophenyl)urea (compound 422; Fig. 6a) as a concentration-dependent growth inhibitor of WT; reducing growth by 14% in MH media at 20 µM (Fig. 6b). This reduction was statistically significant compared to WT grown without compound 422 (*p* = 0.0032; Table S4). The growth of Δ*sbmA* was not significantly affected by the compound at any concentration (Fig. 6c). Next, we performed an LDH assay on macrophages exposed to increasing concentrations of compounds 422. No significant cytotoxicity was observed at the tested concentrations, with p-values indicating no statistical difference from untreated controls (compound 422: 20 µM = 0.0676, 10 µM = 0.1556) (Fig. 6d). We also conducted infection assays using WT *K. pneumoniae* co-cultured with BALB/c macrophages in the presence or absence of compound 422. We observed a statistically significant reduction in CFU recovered from the culture supernatant upon treatment with compound 422 by 31% at 20 µM (p = 0.0047), 22% at 10 µM (p = 0.0600), and 26% at 1 µM (p = 0.0196) (Fig. 6e). Since these compounds only moderately decreased bacterial growth (9-17%), the greater reduction in CFU likely reflects increased phagocytosis. Interestingly, CFU counts from burst macrophages did not differ significantly between untreated WT and compound-treated conditions; however, an increasing trend in CFU from treated compared to untreated *K. pneumoniae* (compound 422: 40% increase, p = 0.6241) was observed (Fig. 6f). These data suggest that the antibacterial effect is dependent upon SbmA and propose an antimicrobial strategy to disrupt bacterial survival through exploitation of nutrient acquisition systems.

**Figure 6:**
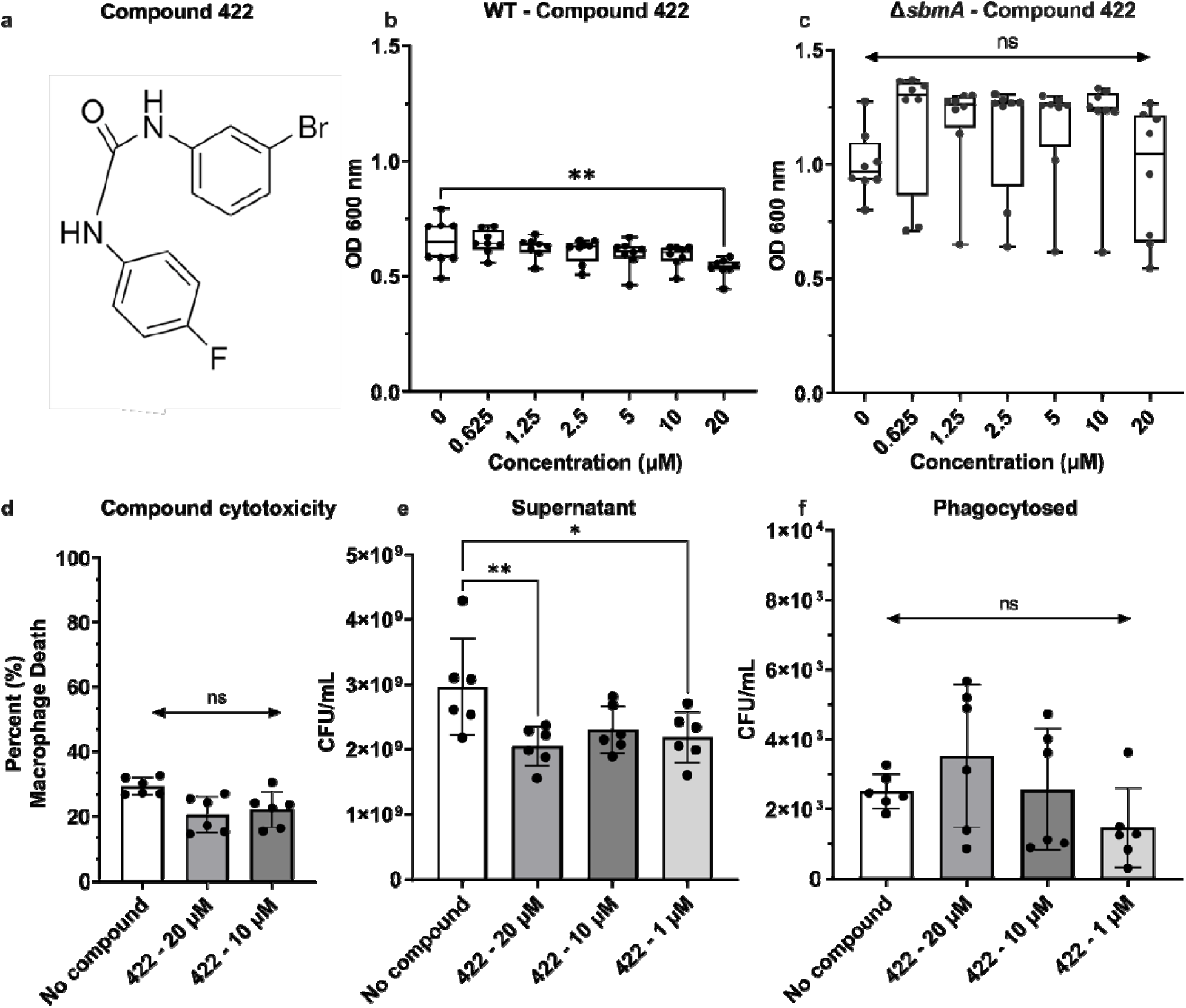
High-throughput drug screening identification of compound that affects WT but not Δ*sbmA K. pneumoniae.* **(a)** Molecular structure of compound 422, 1-(3-bromophenyl)-3-(4-fluorophenyl)urea. **(b)** Box and whisker plot showing median, second and third quartile and minimum and maximum growth values for WT in serially diluted compound 422. **(c)** Box and whisker plot showing median, second and third quartile and minimum and maximum growth values for Δ*sbmA* in serially diluted compound 422. Growth measured at OD_600nm_ in biological quadruplicate and technical duplicate. Statistical significance was determined using a one-way ANOVA. **(d)** LDH assay showing percent macrophage death in the presence of compound 422 compared to untreated. **(e)** WT supernatant CFU following 90 min co-culture with BALB/c macrophages in the presence of increasing concentrations of compound 422. **(f)** BALB/c macrophage phagocytosed WT CFU in the presence of increasing concentrations of compound 422 during 90 min infection. Error bars denote standard deviation. Statistics were performed using One-way ANOVA, where *p < 0.0332, **p < 0.0021, and ***p < 0.0002.

## Discussion

By quantitative phosphoproteome profiling, we report iron-dependent phosphorylation of SbmA and explore functional implications of gene deletion on iron internalization, phenotypic cellular traits, and bacterial virulence. We complemented our phenotypic assessments with global proteome profiling to further validate a role for SbmA in iron acquisition through observed compensatory proteome remodeling in the absence of the gene. Moreover, drug library screening prioritized a compound targeting SbmA, supporting our proposed strategy to exploit the protein’s putative transport function for the disruption of bacterial cell iron homeostasis and delivery of an antimicrobial compound for therapeutic intervention. Further, given the high conservation of SbmA among Gram-negative bacteria, our findings have potential broad implications for antibiotic development, particularly as iron-transport systems have previously proven to be effective antimicrobial targets (38).

The structural consequences of phosphorylation on bacterial nutrient transporters are under-reported, however, evidence from eukaryotic systems demonstrates that phosphorylation can regulate nutrient uptake by inducing conformational changes (94, 95) and by modulating protein-protein or protein-substrate interactions through the addition of negatively charged phosphate groups (96). For *E. coli* SbmA, which shares approximately 86% sequence identity with *K. pneumoniae*, substrate transport of bleomycin and microcin B17 occurs via the PMF, by traversing protons along a ladder of negatively charged glutamate residues (E203, E378, E193, E276, and E287) gated at the cytoplasmic side by Y285 and D288’ (37, 67). Notably, Kp52.145 SbmA shares the same amino acids at the same positions, suggesting substrate transport occurs through the same PMF-driven mechanism. Our predictive modeling indicates that phosphorylation of SbmA at Y285 induces a conformational change that opens the cytoplasmic gate of the transmembrane pore. We hypothesize that phosphorylation promotes proton translocation by altering the conformation of the gate and by introducing an additional negative charge near the glutamate ladder, thereby enhancing the electrostatic attraction of protons. This idea is supported by experimental evidence showing that phosphorylated residues and glutamates can be functionally interchangeable due to their similar size, structure, and charge(97), raising the possibility that phospho-tyrosine may mimic the role of glutamate residues within the proton-conducting ladder. Excitingly, recent evidence in *E. coli* confirms conformational changes of SbmA, which are consistent with mechanism of substrate internalization using the transmembrane proton gradient (98). These findings further support our predicted structural and conformational changes of SbmA in *K. pneumoniae*, which we propose are modulated by phosphorylation. Notably, in *E. coli* mutation of Y285 to alanine (which cannot be phosphorylated), results in slow or halted transport of bleomycin and microcin B17, suggesting that Y285 and phosphorylation at this site, is important for SbmA transport (67).

Throughout our assays, we built evidence for the role of SbmA in iron acquisition; however, the precise contribution of SbmA phosphorylation to iron acquisition remains to be experimentally validated. We propose that SbmA indirectly drives iron uptake by harnessing the PMF to energize a siderophore receptor, analogous to the TonB system (99). This model is consistent with the compensatory over-production of TonB box-containing proteins and siderophore receptors observed in the Δ*sbmA* mutant, and SbmA’s predicted interaction with siderophore biosynthetic pathways (72–76). Alternatively, SbmA may directly transport iron using a PMF-driven mechanism, like its proposed antibacterial peptide uptake mechanism (67). Although most inner membrane iron transporters rely on ATP or GTP (100, 101), rather than the PMF, exceptions, such as MntH (102), demonstrate that PMF-driven metal transport is possible. Interestingly, SbmA contains an ABC transporter domain, like classic inner membrane siderophore receptors, and has been suggested to be an evolutionary link between ABC transporters and PMF-driven transporters (67). Further experimental work is required to define the molecular details of SbmA-mediated iron transport and to determine whether phosphorylation is essential for this process, and if so, whether it acts through conformational modulation, charge redistribution, or altered interactions with other transport components. Notably, SbmA phosphorylation occurs exclusively under iron-limited conditions, suggesting an adaptive regulatory mechanism in which phosphorylation enhances SbmA activity during nutrient stress. These findings underscore the power of mass-spectrometry-based phosphoproteomics for uncovering previously unrecognized regulatory nodes in bacterial nutrient acquisition.

Phenotypic characterization of Δ*sbmA* was performed to identify evidence of nutrient stress arising from impaired cellular iron acquisition. Analyses included electron microscopy, growth curve measurements, and biofilm production assays. Electron microscopy revealed that Δ*sbmA* cells were smaller and exhibited increased filamentation compared to WT. These morphological differences likely represent an adaptive response to nutrient stress, as filamentous growth increases surface area, thereby enhancing access to nutrient-rich environments and facilitating nutrient uptake (103, 104). Similar filamentation has been documented in other *K. pneumoniae* strains lacking genes involved in iron homeostasis, such as *lon* (41). By remaining attached in filaments, Δ*sbmA* cells can facilitate nutrient exchange within the biomass, an adaptive strategy that promotes survival under stress conditions (104). Unexpectantly, Δ*sbmA* exhibited increased growth compared to WT, despite prior evidence indicating that iron availability enhances bacterial growth(105). The increased surface area observed in Δ*sbmA* SEM micrographs provides a plausible explanation for this outcome: filamentation enhances light scattering, thereby elevating OD values (106). Beyond growth, increased surface area also influences biofilm formation by expanding surface binding capacity, which facilitates attachment and biomass development (85). Thus, the enhanced surface area in Δ*sbmA* relative to WT may account for the observed increase in biofilm formation, despite prior evidence linking higher iron availability to increased biofilm development in *K. pneumoniae* (107). Together, these findings suggest that morphological adaptations in Δ*sbmA*, particularly filamentation, override the expected iron-dependent trends, increasing OD values and enabling increased biofilm formation despite impaired iron acquisition.

Given that iron acquisition is an important part of nutritional immunity (13), we anticipated the observed reduced virulence of Δ*sbmA* compared to WT as measured by macrophage LDH release. Studies show that disruption in iron acquisition systems (such as siderophore production) results in reduced virulence in the form of decreased survival within human serum, adhesion and invasion of host cells, and cell death of host cells (108–110). Conversely, hypervirulence has been attributed to increased siderophore production and iron acquisition in bacteria (109, 111). Iron acquisition in bacteria is also associated with increased phagocytic evasion (80). We observed this trend within the present study with macrophage infection assays showing significantly more viable bacteria within Δ*sbmA*-infected macrophages compared to WT-infected. Additionally, in this work, we showed that Δ*sbmA* are smaller, filamentous cells that may be easier to phagocytose than WT *K. pneumoniae*. A third factor could be the broad specificity of SbmA which is known to internalize antibacterial peptides (67). Macrophages, and other immune cells, including neutrophils, use a combination of enzymes, peroxides, low pH and antibacterial peptides to eliminate phagocytosed bacteria (112–117). We hypothesize that the SbmA transporter is exploited by macrophages to internalize antibacterial peptides, which results in much higher clearance rates for WT cells compared to Δ*sbmA* cells within the phagolysosome.

SbmA is recognized as a promiscuous transport protein and, notably, iron transport proteins like SbmA are often exploited by competing bacterial species to deliver toxins that disrupt critical cellular processes (69–71, 118). This susceptibility positions SbmA as a promising candidate for targeted antibacterial development. To identify novel antibacterial compounds that utilize SbmA for entry, we performed a high-throughput screen comparing WT and Δ*sbmA* strains. Compound 422 emerged as a hit, reducing WT growth by 14% while Δ*sbmA* cells showed no growth inhibition under the same conditions. This suggests that the compound’s activity is dependent on SbmA-mediated uptake. Further, no host cytotoxicity and reduced bacterial survival within the supernatant of a macrophage-co-culture system treated with the compound support its potential as a therapeutic. Future studies will assess the efficacy of this compound within *in vivo* infection models along with potential synergistic effects with current antimicrobials targeting *K. pneumoniae*. These findings underscore the value of high-throughput screening in identifying compounds that exploit bacterial transport pathways and target essential cellular functions, accelerating the discovery of new antimicrobial agents.

## Conclusions

In summary, our findings establish SbmA as a previously unrecognized, phosphoregulated contributor to iron homeostasis in *K. pneumoniae*, linking nutrient acquisition with cellular adaptation under iron-limited conditions. By integrating phosphoproteomics with functional and chemical screening, this work not only expands our understanding of bacterial iron regulation but also highlights SbmA as a promising therapeutic target for exploiting vulnerabilities in antimicrobial-resistant pathogens.

## Author Contributions

C.R. & J.G.-M. conceptualized the study. C.R., J.S., & J.G.-M. designed the study. C.R., J.S., S.C & N.C. performed experiments and data analysis. C.R. & O.R. designed and developed figures. C.R. & J.G.-M. wrote and edited the manuscript. All authors have read and approved the submitted manuscript.

## Acknowledgments

Thank you to Rapid Novor for allowing the use of their mass spectrometers, SPARC Drug Discovery at The Hospital for Sick Children for assistance with high-throughput compound screening, and the Molecular and Cellular Imaging Facility and the University of Guelph for assisting with electron microscopy sample preparation and imaging.

## Data Availability

Data are available via ProteomeXchange with identifier PXD072239.

## Supplemental material

Table S1: Primer design for generation and confirmation of *sbmA*Δ.

Table S2: Δ*sbmA* genome missing coverage compared to WT (FO834906.1 accension number), indicating the entirety of *sbmA* is deleted.

Table S3: Receptor proteins significantly more abundant in Δ*sbmA* proteome compared to WT.

Table S4: One-way ANOVA p-values for compound 422 growth inhibition for WT and Δ*sbmA* at different concentrations.

Figure S1: Serially diluted iron standard (Thermo, EEA009) with absorbance measured at 593 nm.

Figure S2: Standard curve for serially diluted (a) WT and (b) *ΔsbmA K. pneumoniae*.

Figure S3: Optical density (OD_600nm_) of WT (a) and Δs*bmA* (b) following incubation with 2,500 drug-like compound library.

Figure S4: Percent of shared sequence identity between bacterial species for SbmA based on BLASTp search.

Figure S5: Superimposition of AlphaFold 3 predicted structure of *K. pneumoniae*-SbmA (A0A2S8TRL6) compared to *E. coli*-SbmA (P0AFY6; PDB accession number 7P34).

