## Supplementary material for "SbmA coordinates iron homeostasis and antimicrobial susceptibility in *Klebsiella pneumoniae* through phosphorylation-dependent regulation": Supp. Files

**Table S1: Primer design for generation and confirmation of ∆*sbmA*.**

| Gene of interest | Forward (F)/ Reverse (R) | Primer Sequence |
| --- | --- | --- |
| *sbmA* + Cm tail | F | GCGTCGTACACAGCAATAGCAGAACGATAAGAAG  TTAGCAGGAGACAATGGTGTAGGCTGGAGCTGCTTC |
| *sbmA* + Cm tail | R | ACGGGAATAATG  CGCGATAAACGTGCGGCCATTATGCTTCCTT  GTCTTGACATATGAATATCCTCCTTAG |
| *sbmA* (size shift) | F | GAAACAATTCTGCACGCATG |
| *sbmA* (size shift) | R | TACCGACGCCACGTCAACAG |
| *sbmA* (in/out) | R | TACCGACGCCACGTCAACAG |
| *Cat* (in/out) | F | GCAACTGACTGAAATGCCTC |

**Table S2: Δ*sbmA* genome missing coverage compared to WT (FO834906.1 accension number), indicating the entirety of *sbmA* is deleted.**

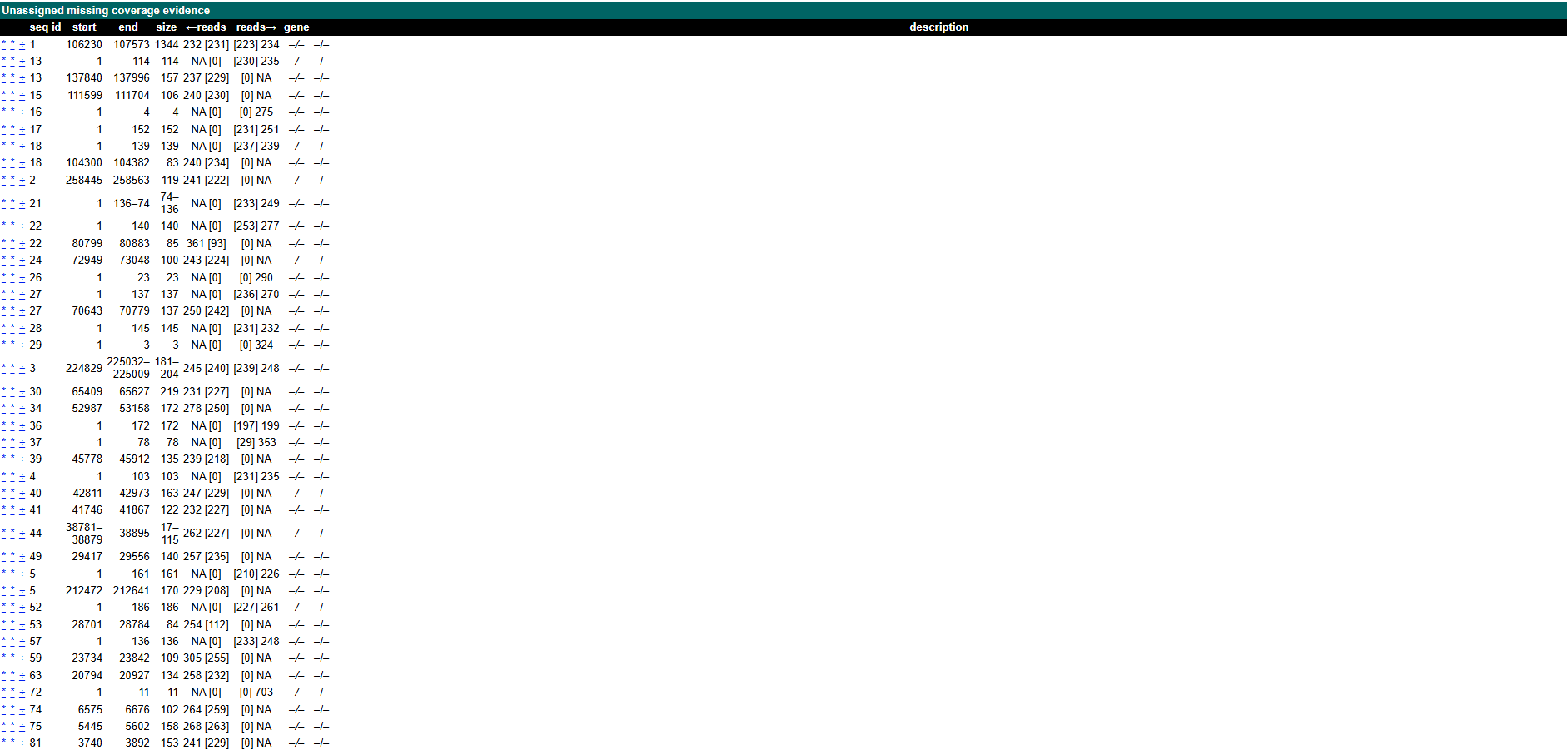

**Table S3: Receptor proteins significantly more abundant in ∆*sbmA* proteome compared to WT**

| **Fold difference** | **UniProt identifier** | **Gene name** | **Protein description** |
| --- | --- | --- | --- |
| 2.87 | A6T7W0 | *yciD* | Outer membrane protein W |
| 3.03 | A6T4V4 | *fhuA* | Outer membrane pore protein, receptor for ferrichrome |
| 3.12 | A6T5E4 | *tsx* | Nucleoside-specific channel-forming protein |
| 3.62 | A6TGF0 | *btuB* | Vitamin B12 transporter |
| 4.29 | A6T631 | *fepA* | Outer membrane porin, receptor for ferric enterobactin and colicins B and D |
| 5.08 | A6T9R9 | *yncD* | Probable tonB-dependent receptor |
| 7.33 | A6TAV9 | KPN_02304 | Ferrioxamine receptor |
| 15.21 | A6T7H0 | KPN_01108 | Ferric aerobactin receptor |

**Table S4: One-way ANOVA p-values for compound 422 growth inhibition for WT and Δ*sbmA* at different concentrations.**

| Strain | Comparison | *p*-value | Significance |
| --- | --- | --- | --- |
| WT | 0 µM vs. 20 µM | 0.0032 | ** |
| WT | 0 µM vs. 10 µM | 0.2452 | ns |
| WT | 0 µM vs. 5 µM | 0.3771 | ns |
| WT | 0 µM vs. 2.5 µM | 0.6845 | ns |
| WT | 0 µM vs. 1.25 µM | 0.9555 | ns |
| WT | 0 µM vs. 0.625 µM | >0.9999 | ns |
| Δ*sbmA* | 0 µM vs. 20 µM | 0.9996 | ns |
| Δ*sbmA* | 0 µM vs. 10 µM | 0.4981 | ns |
| Δ*sbmA* | 0 µM vs. 5 µM | 0.7028 | ns |
| Δ*sbmA* | 0 µM vs. 2.5 µM | 0.8152 | ns |
| Δ*sbmA* | 0 µM vs. 1.25 µM | 0.5627 | ns |
| Δ*sbmA* | 0 µM vs. 0.625 µM | 0.5824 | ns |

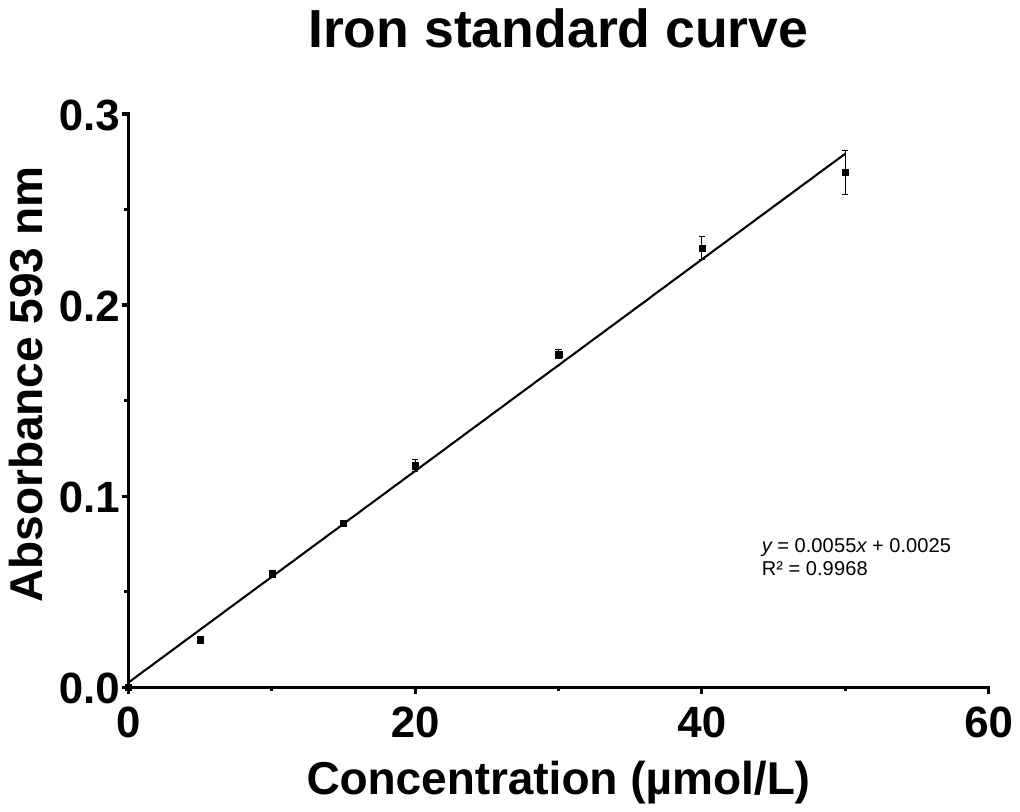

**Figure S1:** Serially diluted iron standard (Thermo, EEA009) with absorbance measured at 593 nm. Experiment performed in duplicate. Error bars denote value range. Curve equation was used to calculate iron concentration per 10^9^ cells for WT and Δ*sbmA* samples.

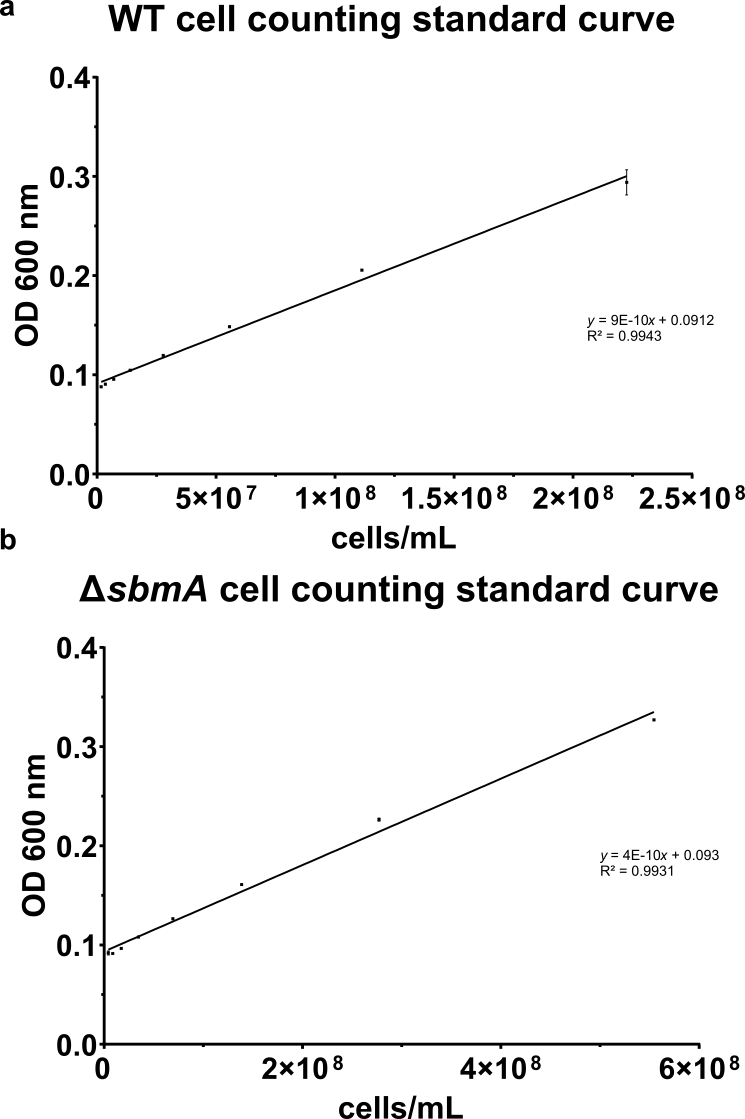

**Figure S2: Standard curve for serially diluted (a) WT and (b) *ΔsbmA K. pneumoniae*.** Absorbance was measured at OD_600nm_ and performed in duplicate. Error bars denote value range; error bars smaller than symbol were not included. Curve equation was used to calculate cells/mL for WT and Δ*sbmA* samples.

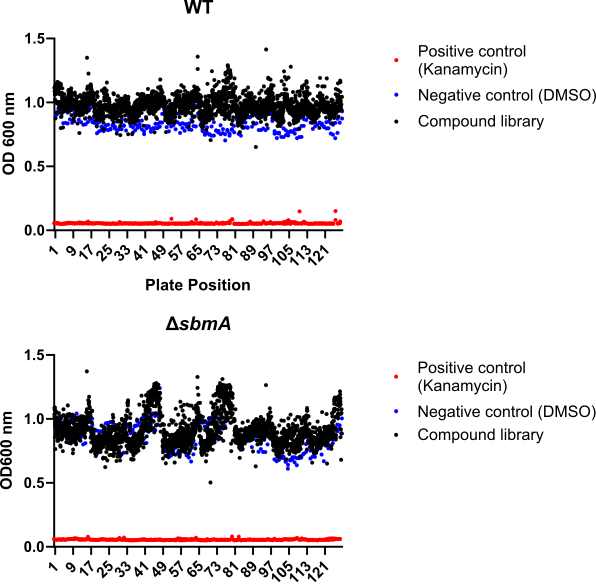

**a**

**b**

**Figure S3: Optical density (OD_600nm_) of WT (a) and Δs*bmA* (b) following incubation with 2,500 drug-like compound library.** Overnight cultures prepared in LB media in 384-well plates containing 10 µM of each compound (dissolved in DMSO). Plates were incubated for 24 h at 37 °C with shaking at 200 rpm. Kanamycin (50 µg/mL) was included as a positive control (red) compound (black) and DMSO (blue) was included as a negative control. Experiment performed in three biological and two technical replicates.

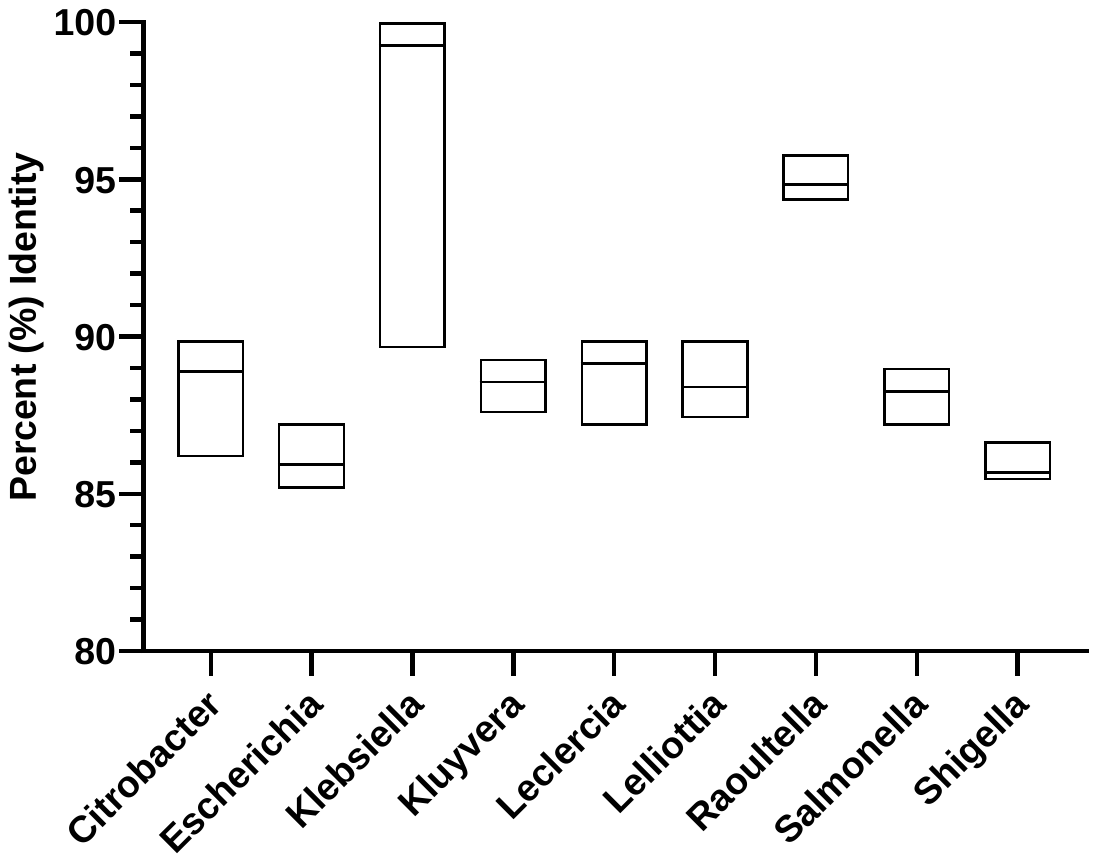

**Figure S4: Percent of shared sequence identity between bacterial species for SbmA based on BLASTp search.** Maximum number of target species was set to 5,000. Floating bars illustrate the median, second and third quartile values for each data set.

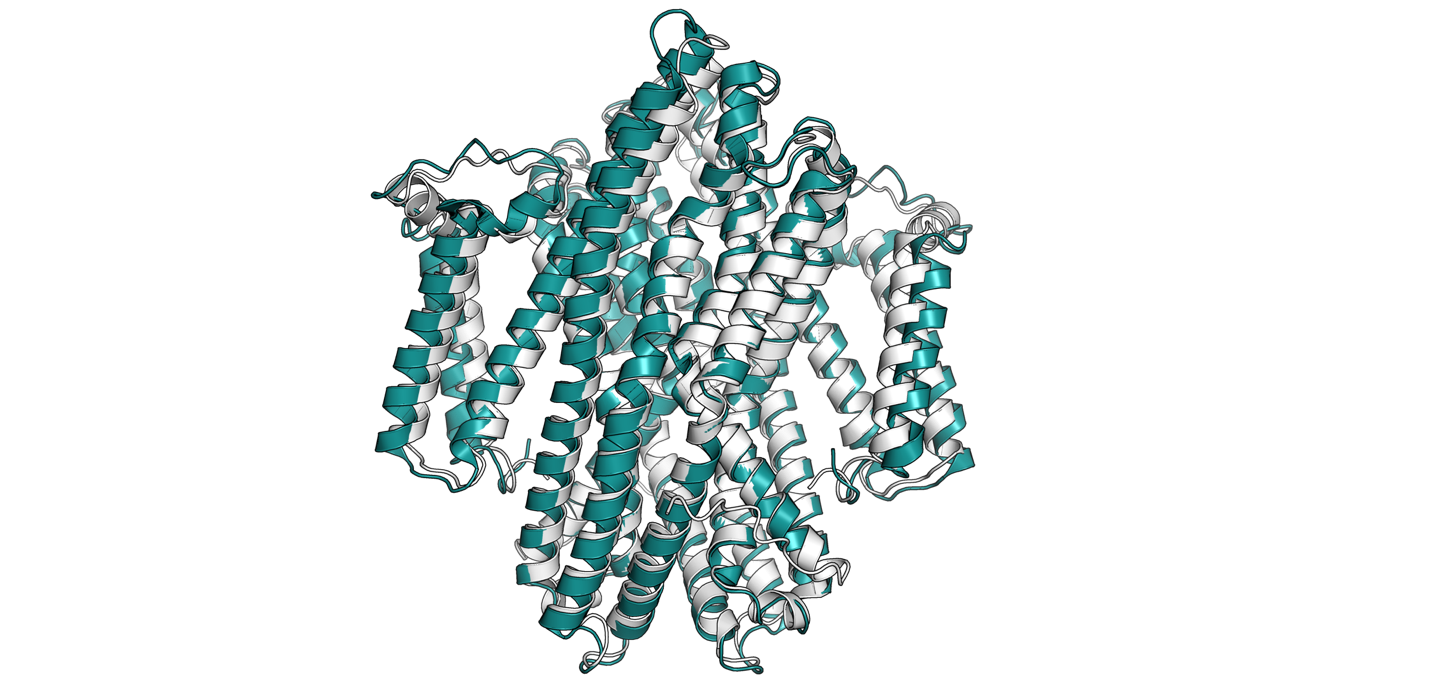

**Figure S5:** **Superimposition of AlphaFold3 predicted structure of *K. pneumoniae*-SbmA (A0A2S8TRL6) compared to *E. coli*-SbmA (P0AFY6; PDB accession number 7P34).** Root mean square deviation (RMSD) of Kp-SbmA/Ab-SbmA was 1.054.
